# Designing combination therapies using multiple optimal controls

**DOI:** 10.1101/850693

**Authors:** Jesse A Sharp, Alexander P Browning, Tarunendu Mapder, Christopher M Baker, Kevin Burrage, Matthew J Simpson

**Affiliations:** School of Mathematical Sciences, Queensland University of Technology (QUT), Australia; ARC Centre of Excellence for Mathematical and Statistical Frontiers, QUT, Australia; Department of Computer Science, University of Oxford, UK (Visiting Professor)

**Keywords:** Leukaemia, Chemotherapy, Stem cells, Combination therapy, Optimal control, Acute myeloid leukaemia (AML)

## Abstract

Strategic management of populations of interacting biological species routinely requires interventions combining multiple treatments or therapies. This is important in key research areas such as ecology, epidemiology, wound healing and oncology. Despite the well developed theory and techniques for determining single optimal controls, there is limited practical guidance supporting implementation of combination therapies. In this work we use optimal control theory to calculate optimal strategies for applying combination therapies to a model of acute myeloid leukaemia. We consider various combinations of continuous and bang-bang (discrete) controls, and we investigate how the control dynamics interact and respond to changes in the weighting and form of the pay-off characterising optimality. We demonstrate that the optimal controls respond non-linearly to treatment strength and control parameters, due to the interactions between species. We discuss challenges in appropriately characterising optimality in a multiple control setting and provide practical guidance for applying multiple optimal controls. Code used in this work to implement multiple optimal controls is available on GitHub.

## 1 Introduction

Determining appropriate interventions for managing populations of interacting species poses significant challenges. A wide variety of biological processes are characterised by interactions between species, ranging from mutualistic to antagonistic. Mutualistic interactions benefit all species involved: for example, the acacia-ant and acacia tree have a mutualistic interaction; the ants are provided food and shelter by the tree, and in turn protect the tree from herbivores, insects and other plants [38]. Antagonistic interactions occur where one species gains at the expense of others, or where all species are disadvantaged; such as the predation of rabbits by foxes [28], competition for prey between lions and hyenas occupying the same ecological niche [81], or in acute myeloid leukaemia (AML) where progenitor blood cells and leukaemic stem cells in the bone marrow niche compete for resources [35,56,80]. These interactions, coupled with the reality that many interventions impact multiple species within an environment, increase the difficulty of designing intervention strategies. We use optimal control techniques to explore the dynamics of multi-species systems subject to multiple intervention strategies. We present a case study considering a combination therapy approach to treatment; analysing a stem cell model of acute myeloid leukaemia (AML). We subject the system to a chemotherapy control that destroys both leukaemic stem cells and progenitor blood cells, and a stem cell transplant control that replenishes progenitor blood cells. This is an informative system to study as the complexity makes it unclear how to best apply these interventions without applying optimal control techniques, and the antagonistic dynamics of AML are representative of many other systems in biology.

Where available intervention mechanisms incur different costs, target different species or have a level of efficacy that varies based on the state of the system, it is reasonable to consider a strategy with multiple interventions applied in combination. We are interested in applying interventions to interacting multi-species processes to influence the outcomes. Example situations where multiple interventions are employed include aerial baiting and animal trapping for invasive species management in ecology [7]; vaccination, rehydration, antibiotics and sanitation for outbreak control in epidemiology [63]; growth hormone and hyperbaric oxygen therapy for wound healing [1]; and chemotherapy and stem cell transplants for cancer treatment in oncology [14].

Interactions between species increase the complexity involved when considering interventions, but can also provide opportunities. Failure to understand the interactions between species can result in unintended effects of intervention. In the Boodaree National Park in south-eastern Australia, intensive fox baiting was implemented to curb population decline of native animals through predation by foxes. This significantly increased the abundance of wallabies, with the unintended consequence of reducing abundance of some tree species [25]. Conversely, understanding interspecies interactions can provide exploitable opportunities in designing interventions. In cancer research, particular genes have been shown to exhibit cancer-promoting functions. There are significant challenges associated with targeting these genes directly, prompting investigation into means of targeting the upstream signalling pathways that activate the genes [76,87]. Understanding the dynamics of interactions between species and the influence of proposed treatments improves our ability to determine effective intervention strategies.

Mathematical models provide a low-cost, low-risk way of exploring the dynamics of biological processes (and valuably, understanding the risks of proposed interventions) [9,21,42,64]. In cancer research mathematical models have been used to explore key processes such as incidence; pathogenesis; tumour growth; metastasis; immune reaction and treatment [16,23,24,44,57,83]. Optimal control techniques are widely applied, not only in mathematical biology broadly [22,50,51,58], but also in cancer therapy [20,36,79] and AML specifically [78].

Likewise, mathematical models have been used to study stem cell dynamics [59,70], including optimal control of cancer via chemotherapy with consideration for bone marrow destruction [29]. Optimal control techniques have been applied to study various combinations of chemotherapy (both broad spectrum and targeted); radiotherapy and anti-angiogenic therapy in the literature [13,47,48,49,66].

Many interacting population dynamics can be explored through studying coupled systems of differential equations. As a starting point, we could consider two species, *S*_1_(*t*) and *S*_2_(*t*) with general growth and decay functions *g*_1_, *g*_2_ and *d*_1_, *d*_2_, respectively. These systems are readily extended to incorporate multiple interventions or controls, say *u* and *v*, that result in some net impacts *c*_1_ and *c*_2_ on each species. Such a model could take the following form, with all variables implicitly functions of time:

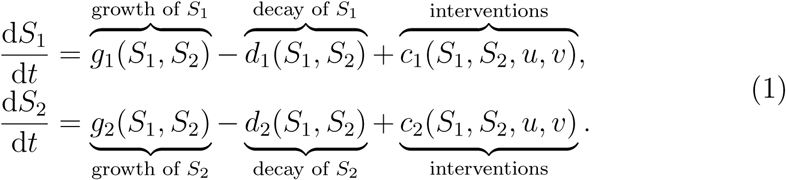

In Equation (1) all growth and decay terms are presented as functions of both species. These terms can be reduced to functions of a single species to capture the range of interactions from mutualism to antagonism. For example, the classic Lotka-Volterra model for predator-prey interaction [11,54] can be recovered if *g*_1_(*S*_1_, *S*_2_) reduces to *g*_1_(*S*_1_), and *d*_2_(*S*_1_, *S*_2_) reduces to *d*_2_(*S*_2_). The intervention terms are expressed as functions of both species and controls for generality, but can also be reduced to model specific interventions. We note that this formulation permits modelling of complex interactions between species and provides flexibility in the application of controls, for example allowing us to implement a control that impacts both species simultaneously.

This paper presents a case study of a two species model with resource competition, where abundance of one species is desirable and the other is undesirable. We consider the dynamics of optimal therapies involving two controls. In particular, we model the dynamics of a combination therapy intervention with one control that negatively affects both species, and another control that positively affects only the desirable species. Results are obtained under pay-off regimes corresponding to continuous applications of both controls; combinations of continuous and bang-bang (discrete) controls; and bang-bang applications of both controls [52,78]. The impact of key parameters on the combination therapy dynamics are also considered. We find that the response of the optimal control strategy to interaction parameters can be highly non-linear, with behaviour that exhibits significant variation across the parameter space. We identify dynamics reflective of clinical practice under some parameter regimes, and note that some interesting solution dynamics are transient, existing only over small regions of control parameter space. We also demonstrate that the response of the system under optimal control dynamics can provide insights into the quality of the underlying model. Finally, we comment on the challenges involved in selecting appropriate pay-off weightings, and the flexibility it affords.

In Section 2 we outline the optimal control approach taken in this work. In Section 3 we introduce a model of AML [23,78] to examine as a case study on combination therapy; we subject the model to both a chemotherapy control and a stem cell transplant control. We identify candidate pay-off functions characterising optimality for the AML model. Results and discussion corresponding to these candidate pay-off functions are presented in Section 4. Concluding remarks are provided in Section 5. In the supplementary material we present a broad collection of results corresponding to a wide range of control parameter regimes.

The code used to produce the optimal control results in this work is freely available on GitHub. Our implementation of the FBSM uses a fourth-order Runge-Kutta method to generate numerical solutions to ordinary differential equations [52,74]. A sufficiently small constant time step is chosen to produce numerical solutions that are grid-independent.

## 2 Optimal control theory

When considering a system with inputs that we can control, we are naturally interested in determining the particular amount and timing of these inputs that produces the *best* outcome. In the context of optimal control, *best* corresponds to a control that minimises or maximises a specified pay-off. The pay-off is also modelled; as such it has assumptions, and in complex systems it is not always clear how to appropriately represent objectives. This is particularly evident when controls are designed to meet multiple objectives; there is not necessarily a single way to value outcomes [60,77]. When considering optimal control for disease treatment, the pay-off typically incorporates factors such as reducing the negative effect of the disease, and minimising the resources used and any adverse effects of the treatment.

For the interacting multi-species system given by Equation (1) a general pay-off functional, *J*, to be minimised over a fixed time interval is:

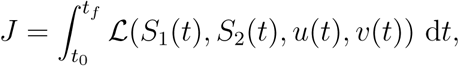

where the integral captures cumulative costs over time, such as disease burden, side effects or toxicity of a treatment, or resource costs. The particular form chosen for ℒ determines the dynamics of the optimal strategy, as we discuss later.

In this work, we determine optimal controls through applications of Pon-tryagin’s Maximum Principle (PMP) [72]. Although multiple interventions are commonly applied in ecology, epidemiology, wound healing and oncology [1,7,14,63], there is limited discussion of optimal control problems with multiple controls. The theory and practice of modelling a single optimal control using the PMP approach has been thoroughly explored in texts such as [3,5,15,52], and extends readily to multiple control problems. As such, we present here only a brief outline of the process, and focus this work on insights and issues of practical implementation of multiple optimal controls. We construct the Hamiltonian and appropriate co-state equations that couple the objective and cost to the multi-species system. Applying the PMP produces a two-point boundary value problem (TPBVP) that we must solve, in combination with known initial conditions for the *state*, {*S*_1_(*t*_0_), *S*_2_(*t*_0_)}, to minimise the Hamiltonian and hence determine the optimal controls and corresponding optimal state trajectories. The TPBVPs arising in optimal control are typically characterised as being a system of differential equations where some initial conditions and some final time conditions are known. As such, they are commonly solved using iterative approaches such as the forward-backward sweep method (FBSM) or shooting methods [40,52].

We apply the FBSM, an iterative process involving the following steps: an initial guess is made for the controls over the interval; using this guess the state is solved forwards in time from *t*_0_ to *t*_*f*_; with this information and the transversality condition (a final time constraint on the co-state variables, derived from the pay-off function), the co-state equations are solved backwards in time from *t*_*f*_ to *t*_0_, and; the guess for the controls are updated based on the solutions for the state and co-state. This process is repeated until the state, co-state and controls are deemed to have converged to some tolerance. Practical guidance and code for implementation of the FBSM for optimal control problems is available in the literature [52,78]. The FBSM readily generalises to problems with multiple controls with no modification beyond including the additional equations, though typically incorporating additional controls also increases the computational cost. An algorithm for the FBSM for multiple controls is provided in the supplementary material.

The pay-off functions we consider vary in regard to whether each of the terms in the integrand are linear or quadratic. Linear and quadratic forms are prevalent in the literature [3,53,68,75], although other forms are also considered, such as a logarithmic pay-off to represent investor utility in mathematical finance [2,45]. Quadratic control terms in the pay-off produce continuous controls. Having the control term appear only linearly in both the pay-off and the state produces bang-bang or singular controls. Bang-bang controls require specified lower and upper bounds, are applied at either bound, and switch based on the sign of the derivative of the Hamiltonian with respect to the control variable. Singular controls arise when this derivative is zero over any finite interval excluding isolated points. Over such intervals the optimal control cannot be determined by simply looking for the value that minimises the Hamiltonian [15,52]. In this work we focus on control parameter regimes where the linear pay-off terms correspond to non-singular bang-bang optimal control problems.

The functional form of state variables in the pay-off has a less clear impact on the control dynamics. Quadratic terms attribute a disproportionally greater cost to large quantities than small; this can be desirable when modelling leukaemia, as a larger leukaemic burden can be disproportionally more damaging than a smaller one [27]. The downside of this is that very little cost is ascribed to a small leukaemic burden, meaning optimal control regimes derived from a pay-off with a quadratic leukaemia term may reach a state where significant leukaemia remains. Conversely, the penalty applied by a linear term is proportional to the size of the leukaemic burden; optimal control regimes derived under this type of pay-off will typically produce final states with less leukaemia remaining.

In the following sections, we explore the dynamics of multiple controls through a model of AML subject to combination therapy. We present select results in this document to highlight key features of multiple control dynamics, and present a broad selection of additional results in the supplementary material.

## 3 Case study: combination therapy for acute myeloid leukaemia

Acute Myeloid Leukaemia (AML) is a blood cancer characterised by the transformation of haematopoietic stem cells into leukaemic blast cells, primarily in the bone marrow [23,65]. The presence of leukaemic cells in the bone marrow niche disrupts haematopoiesis [23], as these cells stop responding to normal regulators of proliferation and no longer undergo normal differentiation or maturation [27,34]. Typical treatments for AML include chemotherapy; immunotherapy; haematopoietic stem cell transplants; radiotherapy and leukapheresis [4,65,73]. Treatment strategies often incorporate multiple therapies concurrently, these combination therapies can be intramodal; such as chemotherapy with multiple chemotherapeutic agents, or intermodal; such as chemotherapy in combination with stem cell transplantation. Combination therapies offer a range of potential advantages over individual therapies, including reduction of toxicity and adverse effects of treatment, improved outcomes in the presence of drug resistance and tumour cell heterogeneity and potentiation of chemotherapy [10,37,85].

In a clinical setting, AML treatment may involve both chemotherapy to reduce the leukaemic cell population and stem cell transplants to bolster the healthy cell populations. This has been shown to allow a higher dose of chemotherapy to be given, reduce adverse effects of the chemotherapy, reduce the risk of relapse and improve long-term survival [14,55,86]. When treating cancer via chemotherapy the exact mechanisms depend on the type of cancer and the particular chemotherapeutic drug, but typically, cytotoxic drugs are administered that target highly proliferative cells. Unfortunately, this commonly includes not only cancer cells, but also healthy cells in the bone marrow, hair, skin and digestive system [65]. The loss of these healthy cells contributes to the significant side effects experienced by chemotherapy patients. A stem cell transplant can mitigate these side effects by giving a patient allogeneic (from a matched donor) or autologous (from the person receiving the transplant) stem cells, typically collected from the bone marrow or the peripheral blood. Transplants are most often administered in remission, following preparative high-dose chemotherapy and/or radiotherapy [14,17]. Side effects arising from stem cell transplants can also be significant, including complications related to the liver, kidneys and lungs, heightened risk of bacterial and viral infections, and graft-versus-host disease [26,32,55].

Development, progression and response to treatment of AML is highly heterogeneous, due to its distinct genetic variation [27,82]. Measuring individual biological parameter values directly from experiments is challenging, and fitting models to clinical data produces parameter estimates that are not unique or with significant uncertainty [33,82]. Further, clinical data is often only collected at a course-grained level, sufficient to describe only the collective behaviour of the system [23,33]. In Ommen et al. (2014), patient data is analysed to investigate the doubling time of the leukaemic burden in relapse of AML [69]. The results are delineated according to molecular subgroups. The median doubling times for these subgroups range from 12 to 24 days. There was significant variance between samples however, with doubling times ranging from 3 days to around 70 days. Another study of patients with untreated, newly diagnosed AML calculates the median potential tumour doubling time to be 8 days, with a range of 3 to 48.9 days [12].

In treating AML with chemotherapy, the timing and dose of chemotherapeutic agents is a critical factor in determining patient outcomes [71]. It follows that the dosages applied in combination therapy are also critical, particularly with studies indicating that synergistic relationships may exist between treatments in particular ratios and schedules but not others [10,61]. As such, it is important to understand the dynamics of systems subject to multiple treatments, and identify key factors that impact the optimal treatment schedules and dosages. Different aspects of treatment schedules interact in a complex way and models are a useful way to investigate the relationship [8].

We consider a stem cell model of AML first presented by Crowell, MacLean and Stumpf [23], modified to incorporate an immune response to leukaemia [78]. The model consists of five species; haematopoietic stem cells *S*(*t*), progenitor cells *A*(*t*), terminally differentiated blood cells *D*(*t*), leukaemic stem cells *L*(*t*), and fully differentiated leukaemia cells *T* (*t*). All dependent variables are functions of *t*, and rates have dimension [*T*^−1^]. We do not explicitly specify a time scale. For notational convenience we use standard variables to denote time dependent quantities:

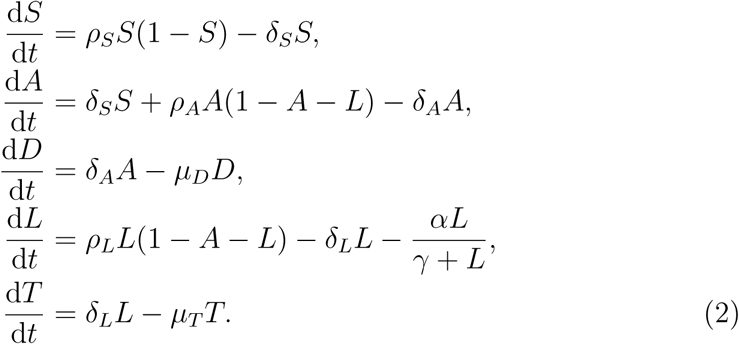

The competition between progenitor blood cells and leukaemic stem cells is based on the hypothesis that these cells occupy the same niche within the bone marrow [35,80]. Motivation for incorporating this kind of interaction in models of AML and other similar leukaemias has been detailed in the literature [23,57].

The steady state behaviour of the original model has been thoroughly explored in [23], and the effect of the incorporated immune response on these steady states is outlined in [78]. Briefly, the original model supports healthy (no leukaemia cells), leukaemic (no healthy cells) and coexisting (both progenitor blood cells and leukaemia cells) steady states, and the immune response incorporated in [78] has the effect of introducing a stable healthy steady state in place of the previously unstable healthy steady state.

The haematopoietic stem cells in the original model grow logistically to steady state: 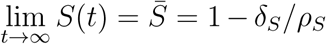, independent of the other cell types. As such, in this work we make a simplifying assumption that the haematopoietic stem cell population is held constant at this steady state. We neglect the terminally differentiated cell populations as they do not feed back to the progenitor or leukaemic stem cell populations, and as such will not influence the design of optimal combination therapy based on progenitor and leukaemic stem cell populations. The chemotherapy control, *u*, is modelled as an additional death term for both progenitor blood cells and leukaemic stem cells. The additional death rate of each type of cell depends on the amount of control applied and the size of the population. The stem cell transplant control, *v*, is modelled as an increase in the progenitor blood cell population, depending only on the amount of control applied. These assumptions reduce Equation (2) to Equation (3). We consider scaled populations such that 0 ≤ *A* + *L* ≤ 1 in absence of control, for suitably chosen initial conditions: *A*(0)+ *L*(0) *<* 1 and *L*(0) ≲ 0.9.

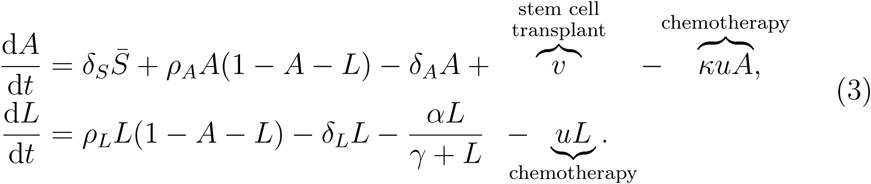

The significant heterogeneity of AML development, progression and response to treatment makes identifying realistic individual parameter values challenging; as such the parameter values in Table 1 are selected such that the qualitative behaviour we observe from the uncontrolled model is reasonable. For illustrative purposes we treat the rates presented to be per day, and present solutions over a potentially typical time-scale of days. We note that there is significant uncertainty in the parameter values, and stress that the focus of this work is on the dynamics and interactions of multiple controls, and in particular, how these dynamics change in response to varying control parameters.

**Table 1:**
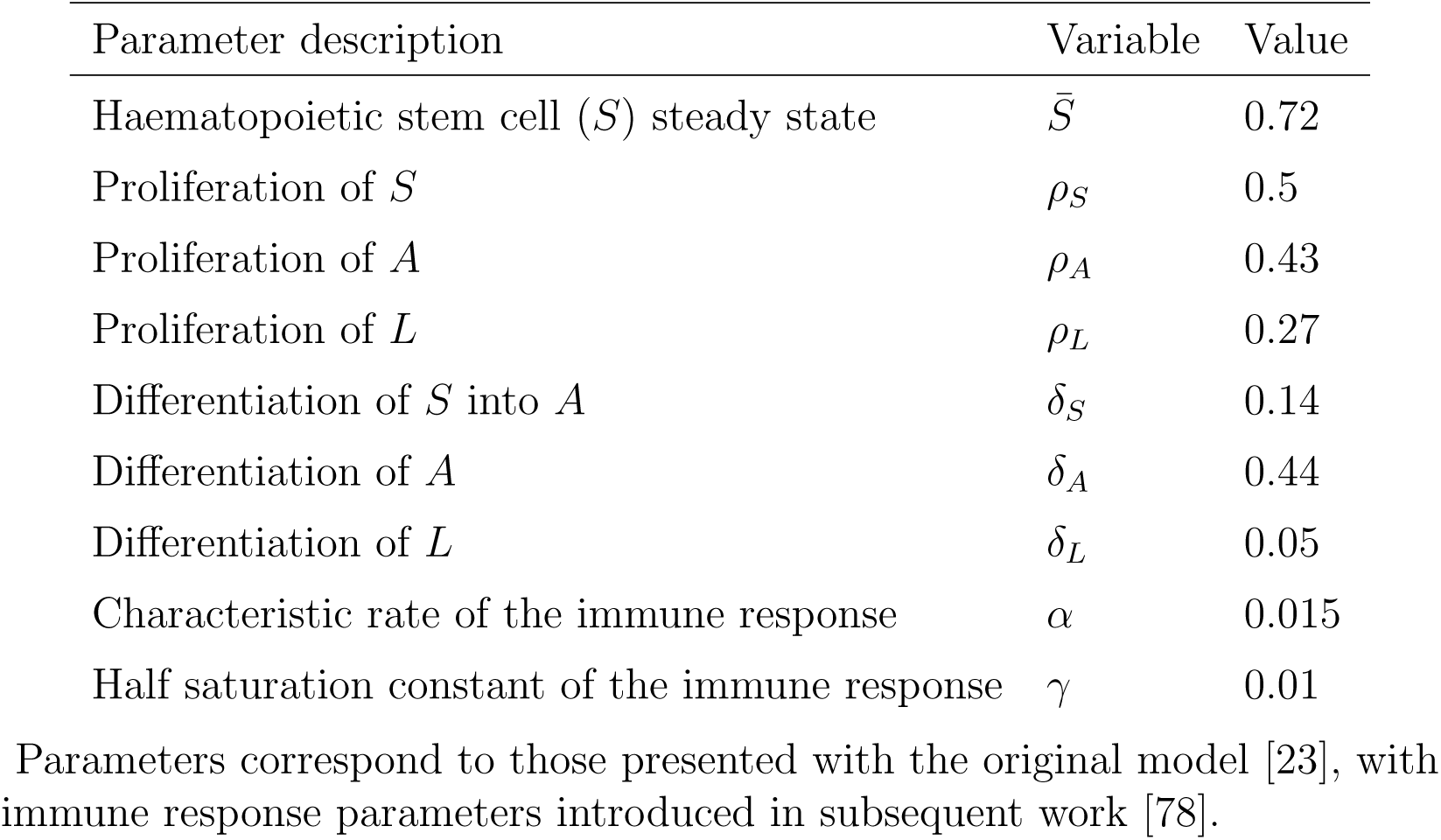
Parameters.

In the context of applying multiple interventions, particularly when an intervention impacts multiple species, understanding interactions between species and controls is crucial for determining appropriate management strategies. As such, we are particularly interested the parameter *κ*, that describes the effectiveness of the chemotherapy in killing progenitor blood cells, relative to leukaemic stem cells; *κ <* 1 corresponds to chemotherapy that is more effective at killing leukaemic stem cells than progenitor blood cells, *κ* = 1 corresponds to chemotherapy that is equally effective at killing either cell type and *κ* > 1 corresponds to chemotherapy that is more effective at killing progenitor blood cells than leukaemic stem cells. This parameter can be adjusted to reflect the clinically observed heterogeneity in response to treatment. A description of the other model parameters and the values used to produce results in this work are presented in Table 1. In absence of control, these parameters can give rise to the coexisting steady state. Dynamics of this model, for various initial conditions, and without any control, are presented in Figure 1.

**Fig. 1.**
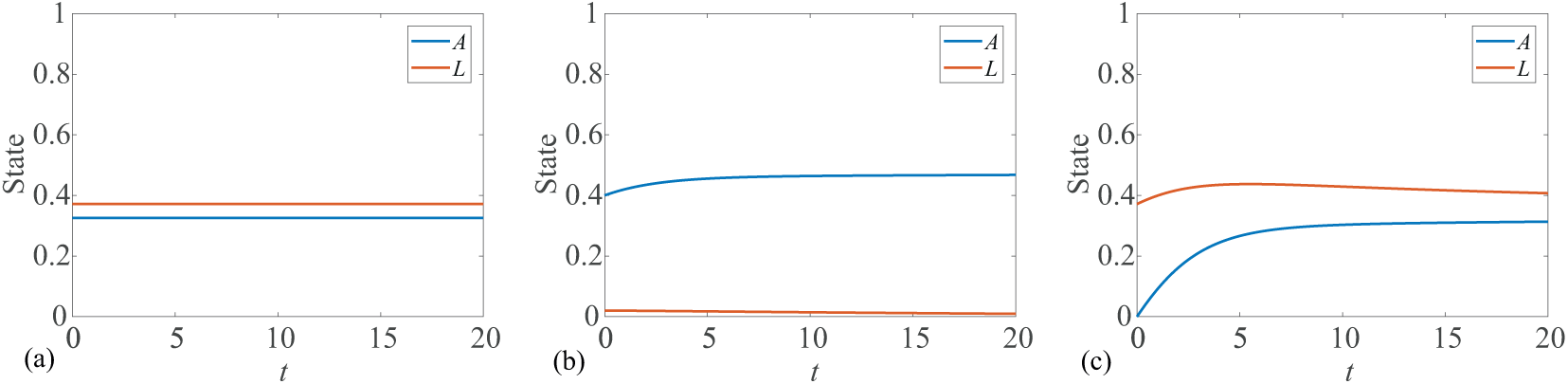
Numerical solutions are presented for Equation (3) to demonstrate the model dynamics for different initial conditions, in absence of control. Initial conditions in (a) correspond to the coexisting steady state. In (b) we observe that a small leukaemic population is depleted through the immune response and competition with progenitor blood cells. In (c) we observe that a small progenitor blood cell population is replenished from haematopoietic stem cells, such that the model tends toward the coexisting steady state.

Due to the significant cost and side effects associated with each treatment, we typically define pay-offs that minimise not only the leukaemic burden but also the amount of each control applied. In the remainder of this section we identify and discuss several reasonable choices of pay-off. We model two controls and explore the dynamics of each possible combination of continuous and bang-bang. This produces four distinct sets of control dynamics. In the supplementary material we present numerical solutions for each combination of control dynamics and investigate the impact of various parameters.

Bang-bang controls require lower and upper bounds on the control variable, and are named as such because the optimal control takes either the lower or upper bound, with finitely many discontinuous switching points throughout the interval. We also apply bounds on the continuous controls. Bounds are used to incorporate practical constraints such as a maximum tolerated dose. In this work, we apply a lower bound of zero to all controls, corresponding to no treatment. The control techniques are general and do not require the lower control bound to be zero. However, this is a practical choice in a treatment context. Unless otherwise specified we choose upper bounds on continuous controls to be sufficiently large; such that they do not constrain the control dynamics. There is no requirement for any relationship between the upper bounds on the chemotherapy and stem cell transplant controls, *u*_*b*_ and *v*_*b*_, respectively.

The pay-offs we consider in this work can be expressed in a general form as:

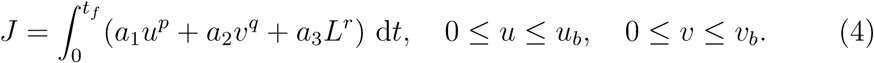

We focus our investigation on the combinations of continuous and bang-bang control possible with *p* ∈ {1, 2} and *q* ∈ {1, 2}. Linear control terms in the pay-off, corresponding to *p* = 1 for chemotherapy control and *q* = 1 for stem cell transplant control, produce bang-bang controls. Quadratic control terms in the pay-off, corresponding to *p* = 2 for chemotherapy control and *q* = 2 for stem cell transplant control, produce continuous controls. In each case we also explore the impact of a linear or quadratic luekaemic term in the pay-off, through choosing *r* ∈ {1, 2}.

Parameters *a*_1_, *a*_2_, *a*_3_ > 0 are chosen to weight the relative importance of each contribution to the pay-off. For example, if we want to assign a greater penalty to the chemotherapy control than to the stem cell transplant control and the leukaemia, we increase *a*_1_ relative to *a*_2_ and *a*_3_. When mixing quadratic and linear control terms care must be exercised when selecting weighting parameters. For the model given by Equation (3), with 0 ≤ *A* + *L* ≤ 1, linear pay-off terms are proportionally more penalising than quadratic terms (*x* > *x*^2^ for *x* ∈ (0, 1)).

### 3.1 Continuous chemotherapy control, continuous stem cell transplant control

For a continuous chemotherapy control and continuous stem cell transplant control we consider a pay-off that minimises the cumulative amount of leukaemia and the controls, with each term squared. This corresponds to Equation (4) with *p* = 2 and *q* = 2.

Results investigating the effect of varying the parameter *κ*; the rate that the chemotherapy control depletes progenitor blood cells relative to leukaemic cells, are presented in Figure S1 of the supplementary material. Results exploring the impact of changing the final time are presented in Figure S2 of the supplementary material.

### 3.2 Continuous chemotherapy control, bang-bang stem cell transplant control

A pay-off that minimises the cumulative amount of chemotherapy control squared, with the stem cell transplant control term entering the pay-off linearly, will produce a continuous chemotherapy control with bang-bang stem cell transplant control. This corresponds to Equation (4) with *p* = 2 and *q* = 1.

Results investigating greater variations in the parameter *κ* are presented in Figure S3 of the supplementary material. Upper bounds on the continuous chemotherapy control are considered in Figure S4 of the supplementary material.

### 3.3 Bang-bang chemotherapy control, continuous stem cell transplant control

Bang-bang chemotherapy control with continuous stem cell transplant control arises from the pay-off in Equation (4) with *p* = 1 and *q* = 2. Noting that each control impacts the state differently (*u* reduces both *A* and *L* while *v* increases *A* only), we can expect this to produce different dynamics to the combination considered in the previous part with *p* = 2 and *q* = 1.

We present results exploring the effect of the final time on the dynamics of the bang-bang chemotherapy control in Figure S5 of the supplementary material. In Figure S6 of the supplementary material we investigate how different upper bounds on the continuous stem cell transplant control impact the dynamics.

### 3.4 Bang-bang chemotherapy control, bang-bang stem cell transplant control

Finally, we investigate the case where both controls enter the pay-off linearly, such that both optimal controls are bang-bang. This corresponds to Equation (4) with *p* = 1 and *q* = 1.

In Figure S7 of the supplementary material, we investigate the impact of increasing the upper bound on each of the bang-bang controls, effectively allowing for stronger doses of each treatment to be applied. In Figure S8 of the supplementary material we consider how the parameter *κ* impacts the dynamics when all controls are bang-bang.

## 4 Results and discussion

In this section we draw insights about the behaviour of the model when subject to interventions, and also more broadly investigate key factors influencing the dynamics of multiple controls. In particular, we focus on the strength of interaction between controls and species, the form and strength of the controls applied, the duration of the treatment interval and the control weighting parameters. Suites of results investigating these aspects are presented in the supplementary material. In the remainder of this section, we highlight key insights.

Due in part to the heterogeneity of AML, there is significant uncertainty around how an individual will respond to treatment [27,43], which is represented by *κ* in our model. This poses challenges in determining appropriate intervention strategies, as it is unclear how heavily chemotherapy treatment will deplete healthy blood cells. As such, a key aspect of this work is to investigate how the optimal control dynamics change as we vary *κ*. Varying *κ* allows us to change the rate that the chemotherapy control depletes progenitor blood cells relative to leukaemic cells. Increasing *κ* makes chemotherapy more damaging to the progenitor blood cells; intuitively one might expect this to promote a reduced application of chemotherapy control. This occurs under some circumstances, but the dynamics are non-linear due to the interactions between the progenitor blood cells and leukaemic cells. In Figure 2 we present results demonstrating that adjusting *κ* can both increase and decrease the duration of both chemotherapy and stem cell transplant controls, depending on the control weighting parameters. Changing control weighting parameters could be thought of as representing the way that different individuals may be more or less heavily impacted by the (side) effects of leukaemia or the treatments [66]. These results agree with the clinically observed heterogeneity in response to regimented treatment, and highlights a key challenge in managing the interacting populations. Even when the nature of an interaction is known, the optimal intervention strategy can vary significantly with changes to the strength of the relationship, the strength of the treatment and the form of the controls.

**Fig. 2.**
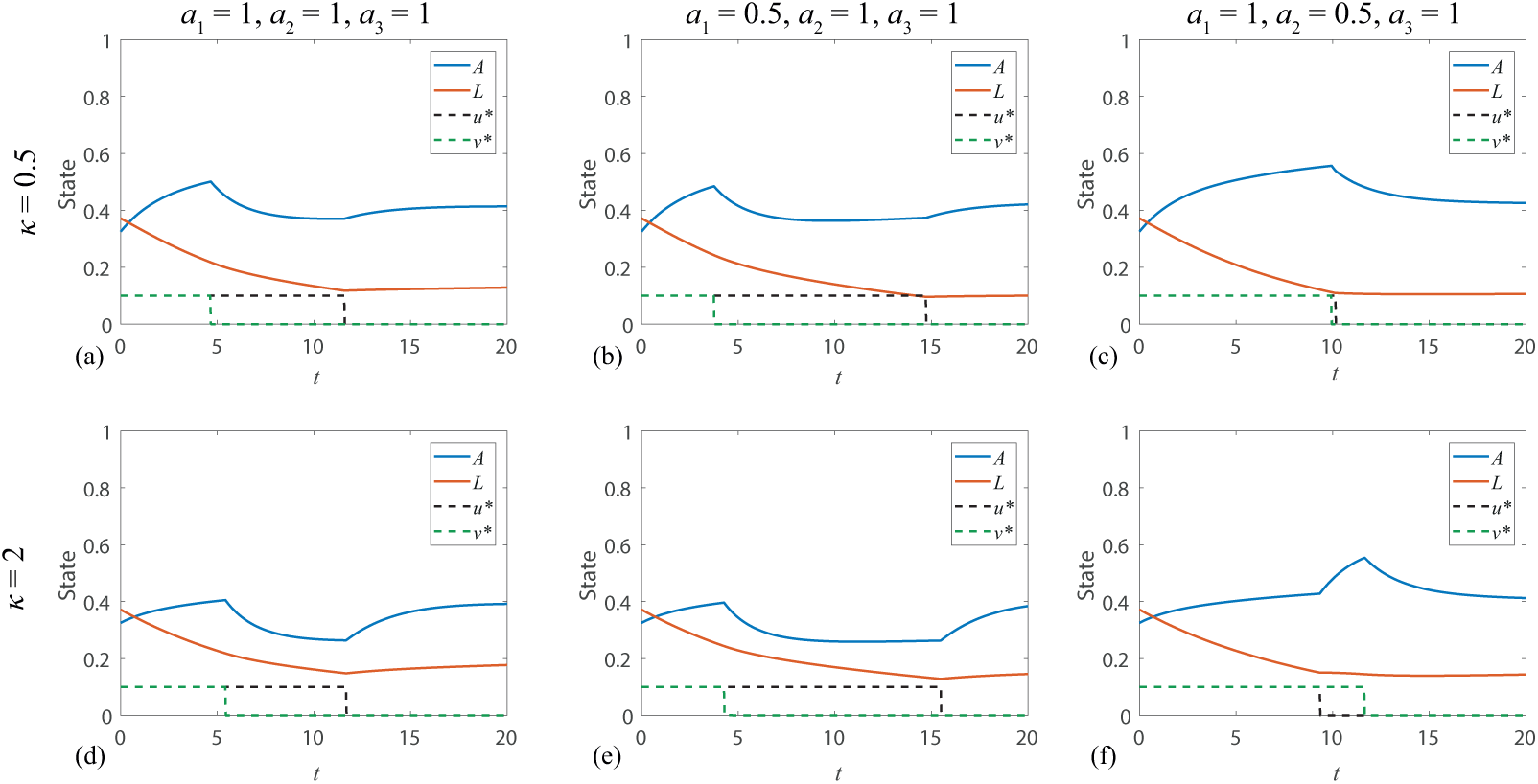
The control dynamics respond non-linearly to the parameter *κ*. Solutions are presented corresponding to the linear pay-off given in Equation (4) with *p* = 1, *q* = 1, *r* = 1 and upper control bounds *u*_*b*_ = *v*_*b*_ = 0.1. With equal weightings (*a*_1_ = *a*_2_ = *a*_3_ = 1), increasing *κ* from *κ* = 0.5 to *κ* = 2 extends the duration of the stem cell transplant control and has little effect on the chemotherapy control. With a reduced weight on the chemotherapy control in the pay-off (*a*_1_ = 0.5), increasing *κ* from *κ* = 0.5 to *κ* = 2 extends the duration of both controls. With a reduced weight on the stem cell transplant control in the pay-off (*a*_2_ = 0.5), increasing *κ* from *κ* = 0.5 to *κ* = 2 increases the application of the stem cell transplant control and reduces the application of the chemotherapy control.

The upper bound on a control represents the maximum strength of the treatment, and can be used to enforce a practical constraint such as a maximum tolerated dose. For bang-bang controls with a lower bound of zero, the optimal treatment is to apply the drug at the maximum tolerated dose (over one or many intervals) or not at all. Bounds can also be used to enforce a dosage threshold on continuous optimal controls, while still admitting intermediate doses. In Figure 3 we observe that increasing the treatment strength results in a reduced duration of application of both controls. For sufficiently high maximum doses only chemotherapy control is applied. At lower *κ*, we observe that the maximum doses must be higher before combination therapy is abandoned in favour of solely chemotherapy. This demonstrates that the strength of a treatment can influence whether or not it is applied, indicating a non-linear response of the optimal control strategy to treatment strength.

**Fig. 3.**
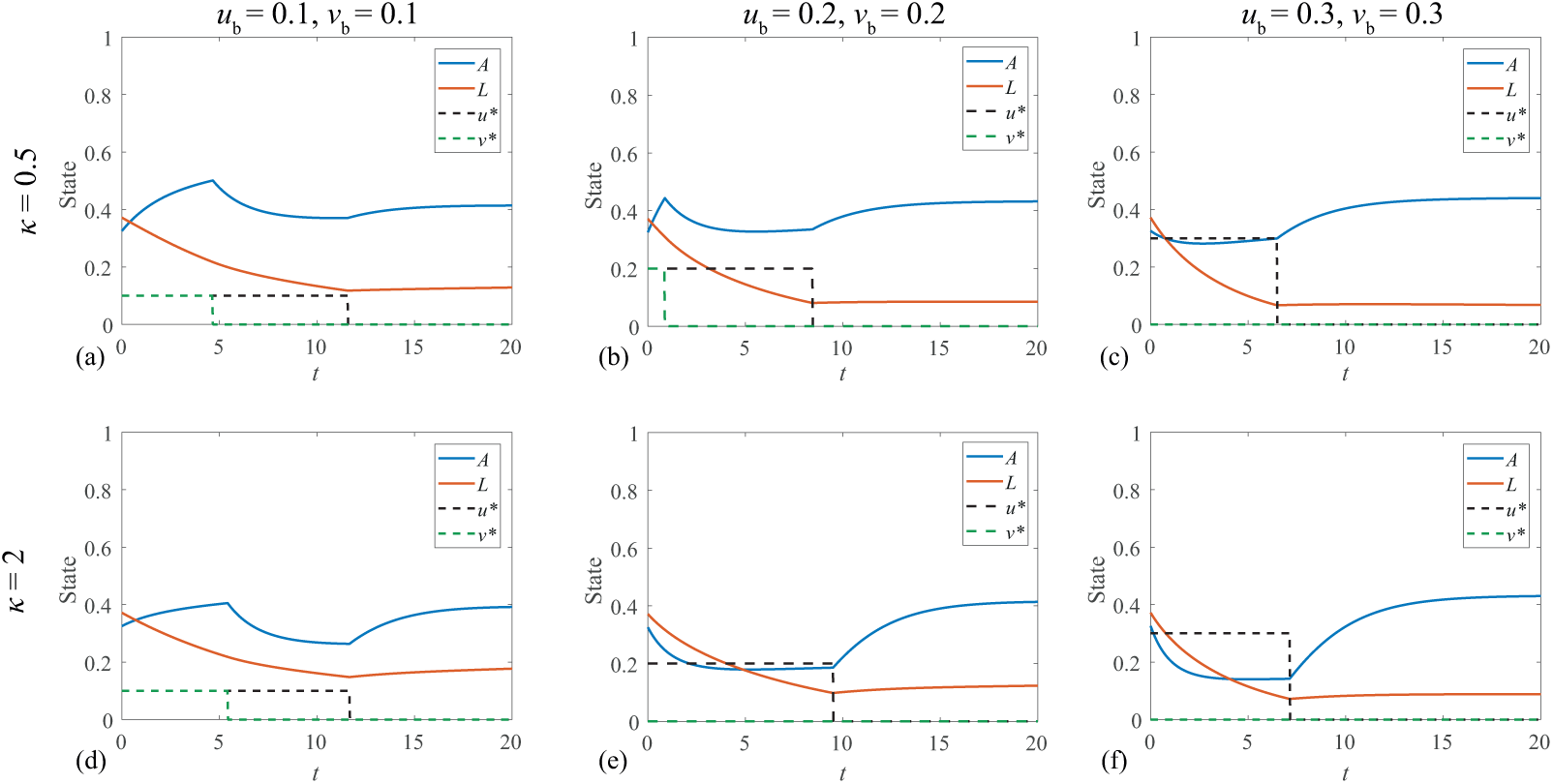
Increasing the upper bound of the bang-bang controls can significantly alter the dynamics. Solutions are presented corresponding to the linear pay-off given in Equation 4 with *p* = 1, *q* = 1, *r* = 1 and equal pay-off weightings (*a*_1_ = *a*_2_ = *a*_3_ = 1). When the chemotherapy control is more effective at killing leukaemia cells than progenitor blood cells (*κ* = 0.5), we observe that increasing the upper bound of each control from *u*_*b*_ = *v*_*b*_ = 0.1 to *u*_*b*_ = *v*_*b*_ = 0.2 results in a reduced duration of application of both controls. Increasing the bounds further to *u*_*b*_ = *v*_*b*_ = 0.3 leads to the result in panel (c) where only the chemotherapy control is applied. When the chemotherapy control is more effective at killing progenitor blood cells than leukaemia cells (*κ* = 2), the stem cell transplant control is no longer applied after increasing the upper bound of each control from *u*_*b*_ = *v*_*b*_ = 0.1 to *u*_*b*_ = *v*_*b*_ = 0.2.

Optimal control dynamics that resemble clinical practice are recovered under particular pay-off weighting parameters. If a stem cell transplant is administered in practice, it typically follows high-dose chemotherapy or radiotherapy to reduce the level of leukaemic cells, suppress the immune system and condition the patient for the introduction and growth of the new blood cells [18,62]. In Figure 4(b) we observe that for particular pay-off weightings it is possible to recover similar behaviour through optimal control solutions to the model.

**Fig. 4.**
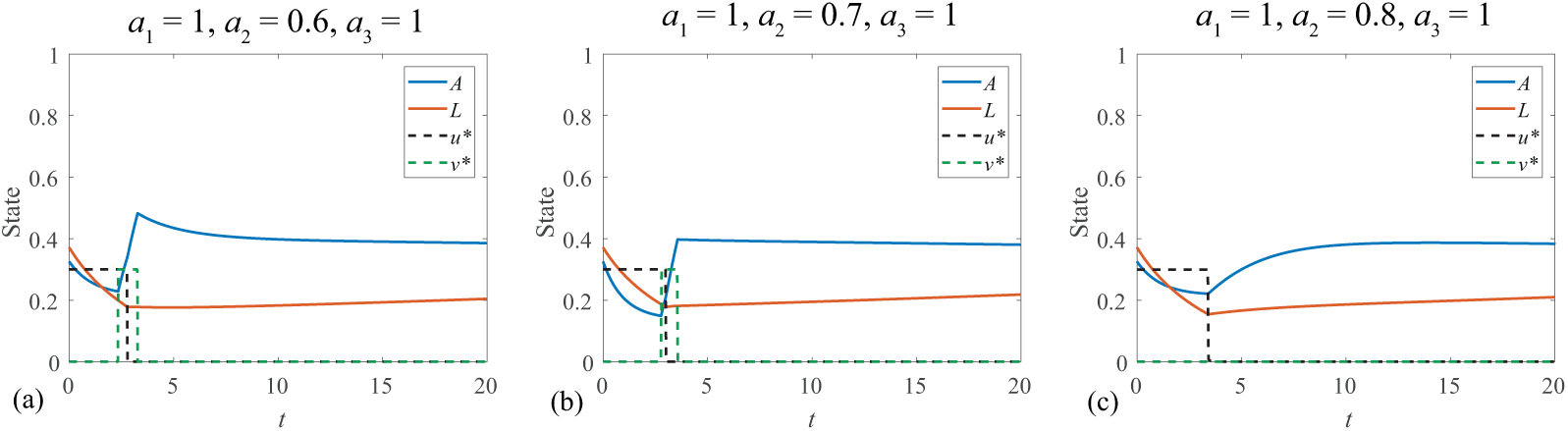
Under particular pay-off weightings, we observe that the stem cell transplant control is applied as the chemotherapy treatment is stopped. Solutions are presented corresponding to the pay-off given in Equation 4 with *p* = 1, *q* = 1, *r* = 2 and upper control bounds *u*_*b*_ = *v*_*b*_ = 0.3.

It is also interesting to note the transience of this result within the pay-off weighting parameter space. In Figure 4(a) a small decrease in the weighting on the stem cell transplant control in the pay-off causes the stem cell transplant control to be applied earlier and for longer. Conversely, the small increase in the weighting on the stem cell transplant control demonstrated in Figure 4(c) results in no stem cell transplant control being applied at all. However, under these parameters the optimal control results exhibit a significant leukaemia population remaining at final time.

Optimal control results can also provide insight into the quality of the underlying model and its assumptions. In this work we consider a simplified stem cell model of acute myeloid leukaemia. In reducing this model, it is assumed that the haematopoietic stem cell population is held constant at its steady state, allowing us to focus on the dynamics of the progenitor blood cells (that are replenished at a constant rate due to the haematopoietic stem cell population) and the leukaemic cells. This assumption is reasonable for a large region of control parameter space, although it is quite restrictive when we are considering control regimes that would significantly deplete the progenitor population, such as where *κ* and *u*_*b*_ are large. This can be observed in Figure 3(e) where the progenitor blood cell population remains almost constant from *t* ≈ 5 to *t* ≈ 10 despite the chemotherapy control being applied up to *t* ≈ 10, due to the constant rate of replenishment of progenitor blood cells. This causes the stem cell transplant control to be devalued in these circumstances, suggesting that the model could be improved by reintroducing the haematopoietic stem cell population, and having it also negatively impacted by the chemotherapy control.

It is clear that the optimal control strategy entirely depends on the form and weighting of the pay-off. The results in the main text and the associated supplementary material demonstrate that these choices can produce fundamentally different optimal control dynamics and outcomes. Selecting an appropriate pay-off form and weighting is therefore a significant and recurring challenge in applied optimal control. An ideal pay-off encodes how each factor is truly valued relative to each other factor, although determining this is often not feasible or requires subjective judgement. Determining appropriate pay-off functions becomes increasingly complicated when dealing with multiple controls; it introduces the need for relative weighting, and in some cases scaling or normalisation of pay-off terms. Control problems in biology typically include non-monetary costs; such as side-effects of treatment, which are not easily valued [78,84]. Even for problems where all elements of a pay-off are readily expressed in monetary terms, there can be challenges associated with economic uncertainty [46].

When determining the form of a control term in the pay-off function it may be tempting to consider how the control can be applied practically. For example, if it is only possible for the control to be active or not active (with no intermediate levels of activation) using a linear term ensures that the optimal control is bang-bang. Particular care must be taken in employing this approach for a multi-objective pay-off, as a linear term implies a different cost weighting to (for example) a quadratic term, which penalises disproportionally; (*x* > *x*^2^ for *x* ∈ (0, 1), *x < x*^2^ for *x* ∈ (1, ∞)). A standard approach in the literature is to consider a practical range of parameter values, often guided by the literature or expert opinion, and examine how sensitive the control strategy is to changes in these parameters [39,67]. The parameters can also be tuned to produce optimal control strategies that satisfy additional external constraints [6]. An alternative approach feasible for some problems is to construct the simplest possible pay-off (for example, the sole objective of minimising leukaemia), at the cost of requiring a more sophisticated model that explicitly accounts for the negative impacts or costs of the controls.

In some regions of the parameter space, the FBSM fails to converge. Convergence can often be achieved by adjusting the parameter that weights the contribution of the control from previous iteration to the control in the next iteration [78]. However, there are some regions of the parameter space where the control does not converge for any value of the weighting parameter. Interestingly, this non-convergence does not always occur at the extremities of a parameter space. For example, comparing Figure S3(h) with Figure S3(i) of the supplementary material, we show that increasing the rate that the chemotherapy control kills progenitor blood cells relative to leukaemic cells from *κ* = 1 to *κ* = 10 moves the optimal control solution from a moderate application of both controls, to a solution dominated by stem cell transplant control. For intermediate values, 5.5 ≲ *κ* ≲ 7.5, this control problem does not converge. This may correspond to a region where the optimal control is not bang-bang, but rather is singular. Singular control problems arise when the first order conditions derived from the PMP do not provide sufficient information over an interval to determine the optimal control [15]. Singular controls can sometimes be determined from higher order optimality conditions, although this is non-trivial [30,31].

## 5 Conclusion

In this work we study the application of multiple optimal controls to a stem cell model of AML. We consider a chemotherapy control that reduces the population of both progenitor blood cells and leukaemic stem cells, and a stem cell transplant control that replenishes progenitor blood cells. To investigate the dynamics arising from different forms of control and interspecies interaction, we generate results corresponding to each of the possible combinations of continuous and bang-bang controls, with pay-off functions containing linear or quadratic leukaemia terms. The dynamics of multiple controls are further explored through varying control parameters, treatment strengths and the rate at which the chemotherapy control depletes progenitor blood cells relative to leukaemic stem cells.

We determine optimal controls through application of PMP, yielding two-point boundary value problems. Numerical solutions to these problems are generated using an implementation of the forward-backward sweep method. The method readily generalises to solving problems with multiple controls, and we observe only a modest increase in computational resources required beyond a comparable single optimal control problem.

Through investigating how optimal control dynamics change in response to the relative effectiveness of the chemotherapy control, the maximum strength of the controls and other control and weighting parameters we show that the behaviour can be highly non-linear. We observe that interesting and clinically reflective optimal control strategies can be transient, existing only over small regions of control parameter space. We also demonstrate how optimal control results can provide insight into the quality of the underlying model.

Modelling multiple interventions that incur costs naturally increases the complexity involved in determining appropriate pay-off functions. A pragmatic approach is to consider a range of weighting parameters and observe the optimal control dynamics. The sensitivity of these dynamics to the weighting parameters can provide insight into how carefully a pay-off must be chosen. If the optimal control dynamics under a particular set of control parameters do not represent a desirable outcome to the practitioner, then the pay-off function may need to be modified [41].

The primary avenues for extending this work are model refinement and control approach. The model could be extended to incorporating a delay to the impact of the stem cell transplant on the system, either through a delay differential equation, or an additional state equation acting as a reservoir of *A* cells produced by the stem cell transplant. This is biologically motivated as it can take multiple weeks for the production of blood cells to occur following a transplant [19]. This extension would also facilitate relaxing the assumption of a constant reservoir of haematopoietic stem cells replenishing the stem cell population. The control approach in this work focuses on fixed terminal time problems. Although informative, this can result in optimal control results that are clinically undesirable under some parameter regimes, such as having a significant amount of leukaemia remaining at the terminal time. Alternatively, optimal controls can be determined for problems with free terminal times and fixed final states; for example no leukaemia remaining [52]. The problems explored in this work could be recast as fixed final state problems to determine how significantly this alters the optimal control dynamics.

## 6 Acknowledgements

This work is supported by the Air Force Office of Scientific Research (BAA-AFRL-AFOSR-2016-0007) and the Australian Research Council (DP170100474). Computational resources were provided by the eResearch Office at QUT. J.A.S gratefully acknowledges support from the Australian Government Research Training Program and the AF Pillow Applied Mathematics Trust.

