## Supplementy material document for "Designing combination therapies using multiple optimal controls"

**Contents**

1 Supplementary results 2

1.1 Continuous chemotherapy control, continuous stem cell transplant control 2

1.2 Continuous chemotherapy control, bang-bang stem cell transplant control 7

1.3 Bang-bang chemotherapy control, continuous stem cell transplant control 11

1.4 Bang-bang chemotherapy control, bang-bang stem cell transplant control 15

2 Supporting code 19

3 Forward backward sweep method 19

References 20

* Corresponding author

### 1 Supplementary results

Key results are presented in the Main document to highlight interesting dynamics and identify important parameters. In this Supplementary Material document we present a broad suite of results corresponding to a wider range of parameter values. This section is divided into four subsections, with each subsection corresponding to one of the four possible combinations of control dynamics (continuous and/or bang-bang). Within each subsection, two sets of results are presented; the leukaemic term is included in the pay-off quadratically in the first set and linearly in the second set. To enable comparison, the form of the leukaemic term in the pay-off is the only difference between the central columns (sub-figures (b), (e), (h) and (k)) of figures within a subsection. Rows in each figure correspond to a different combination of pay-off weighting parameters ($a_1$,$a_2$,$a_3$). We explore the impact of varying a specified parameter across the rows of each figure.

#### *1.1 Continuous chemotherapy control, continuous stem cell transplant control*

This Subsection contains results for pay-offs presented in Equation (S1) and Equation (S2), where both controls are continuous. These pay-offs correspond to results in Figure S1 and Figure S2, respectively. The pay-off functions are given by:

$$J = \int_0^{t_f} (a_1 u^2 + a_2 v^2 + a_3 L^2)\ \mathrm{d}t, \tag{S1}$$

$$J = \int_0^{t_f} (a_1 u^2 + a_2 v^2 + a_3 L)\ \mathrm{d}t. \tag{S2}$$

There are a number of intuitive results that we generally observe across all control problems considered in this document. For continuous controls, increasing the pay-off weighting of a control reduces the amount of that control applied, and typically increases the amount of the other control applied. Increasing the weighting of leukaemia in the pay-off typically leads to a greater amount of both controls applied. With bang-bang controls the upper bound or maximum dose is fixed, so increasing the pay-off weighting causes the control to be applied for a shorter duration, or not at all. Increasing the weighting on leukaemia in the pay-off will typically increase the duration over which the bang-bang controls are applied. Incorporating leukaemia in the pay-off quadratically results in a higher level of leukaemia remaining at the terminal time. This is because linear pay-off terms are proportionally more penalising than quadratic terms; making a greater contribution to the pay-off that we are minimising ($L > L^2$ for $L \in (0,1)$). Broadly, the nature of interactions between the controls and the state variables results in the chemotherapy control being applied to reduce the leukaemic population, and competition between progenitor blood cells, bolstered by the stem cell transplant control, and leukaemic cells prevents the leukaemic population from resurging as the chemotherapy control is reduced.

Results in Figure S1 explore the impact of $\kappa$, the parameter that determines the effectiveness of the chemotherapy at killing the progenitor blood cells relative to leukaemic cells. Comparing Figure S1(a) with Figure S1(c), we see that $\kappa$ does not appear to have a significant impact on the control dynamics. Despite the early-time decline of progenitor blood cells we observe in Figure S1(c) in response to the chemotherapy control, at terminal time the remaining progenitor blood cell and leukaemic populations are not significantly different than in Figure S1(a). These results are consistent for the alternative pay-off weightings considered. When leukaemia is weighted more heavily in the pay-off as in Figure S1(j-l), a higher level of chemotherapy control is applied initially, although the state at terminal time does not vary significantly with $\kappa$.

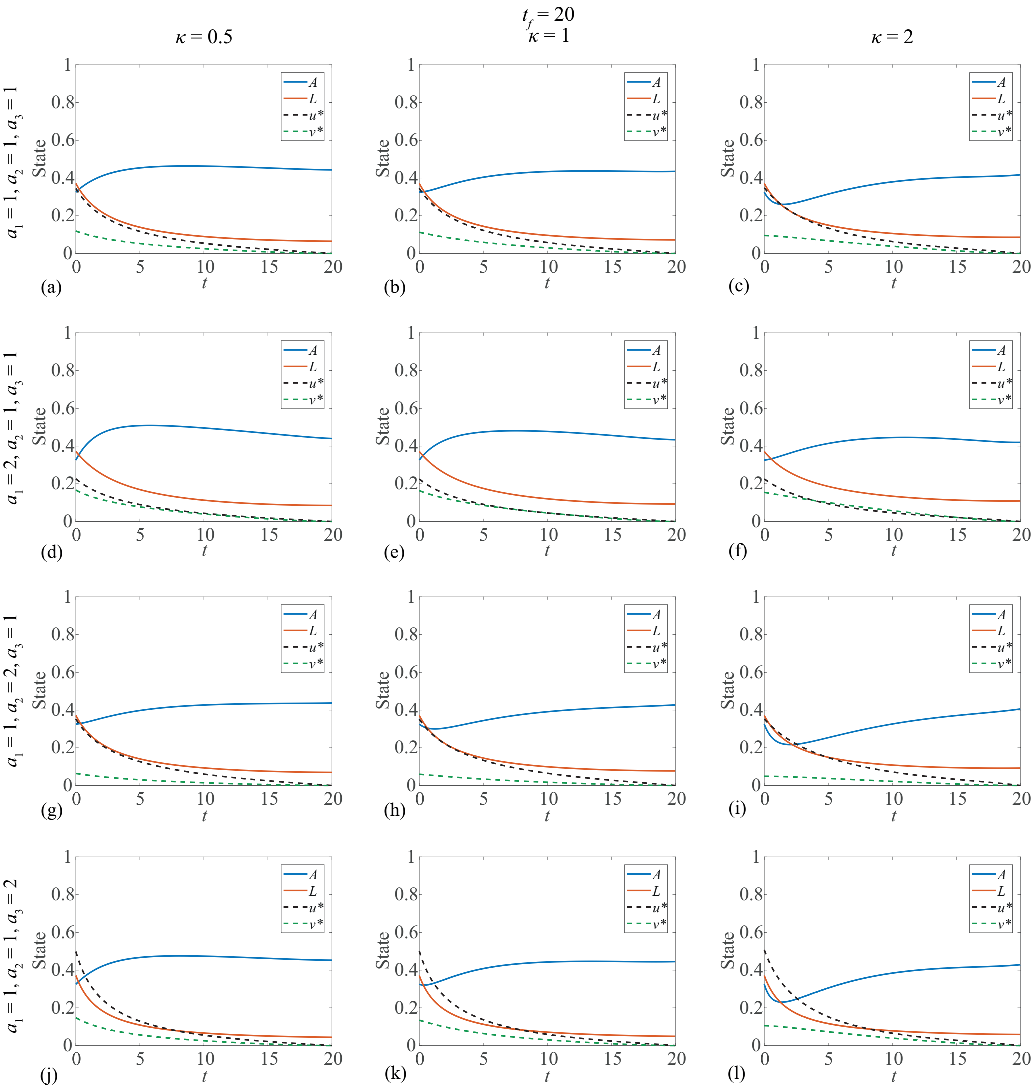


Fig. S1. Results are presented where both controls are applied continuously, under a range of pay-off weightings ($a_1, a_2, a_3 \in \{1, 2\}$) and where the chemotherapy control $u$ affects the progenitor blood cell population less, equally and more than the leukaemic cell population; $\kappa \in \{0.5, 1, 2\}$, respectively. This figure corresponds to the pay-off given by Equation (S1), where the leukaemic term enters the pay-off quadratically. This corresponds to Equation (4) in the Main document with $p = 2$, $q = 2$, $r = 2$.

In Figure S2 we consider the pay-off given by Equation (S2), with the leukaemic term entering the pay-off linearly. Each column corresponds to a different terminal time, to investigate how this impacts the state and control dynamics. Although the results for different terminal times are not a direct scaling of one another, the dynamics are very similar. The results with larger terminal times exhibit a marginally higher initial level of control applied. We also observe a reduced leukaemic population at terminal time when the terminal time is increased. Since the pay-off considers the cumulative leukaemic burden, increasing the terminal time increases the interval over which the leukaemic population contributes to the pay-off, providing a stronger impetus to reduce it.

Comparing Figure S2(b,e,h,k) with Figure S1(b,e,h,k), respectively, we can investigate the impact of incorporating leukaemia in the pay-off linearly rather than quadratically. With linear weighting, leukaemia makes a greater contribution to the pay-off and motivates increased application of control. As such, a significantly higher dose of chemotherapy is applied, to more rapidly reduce the leukaemic population. Corollary to this, a linear leukaemic term results in a lower leukaemic population at terminal time. A modest increase in the stem cell transplant control is also observed.

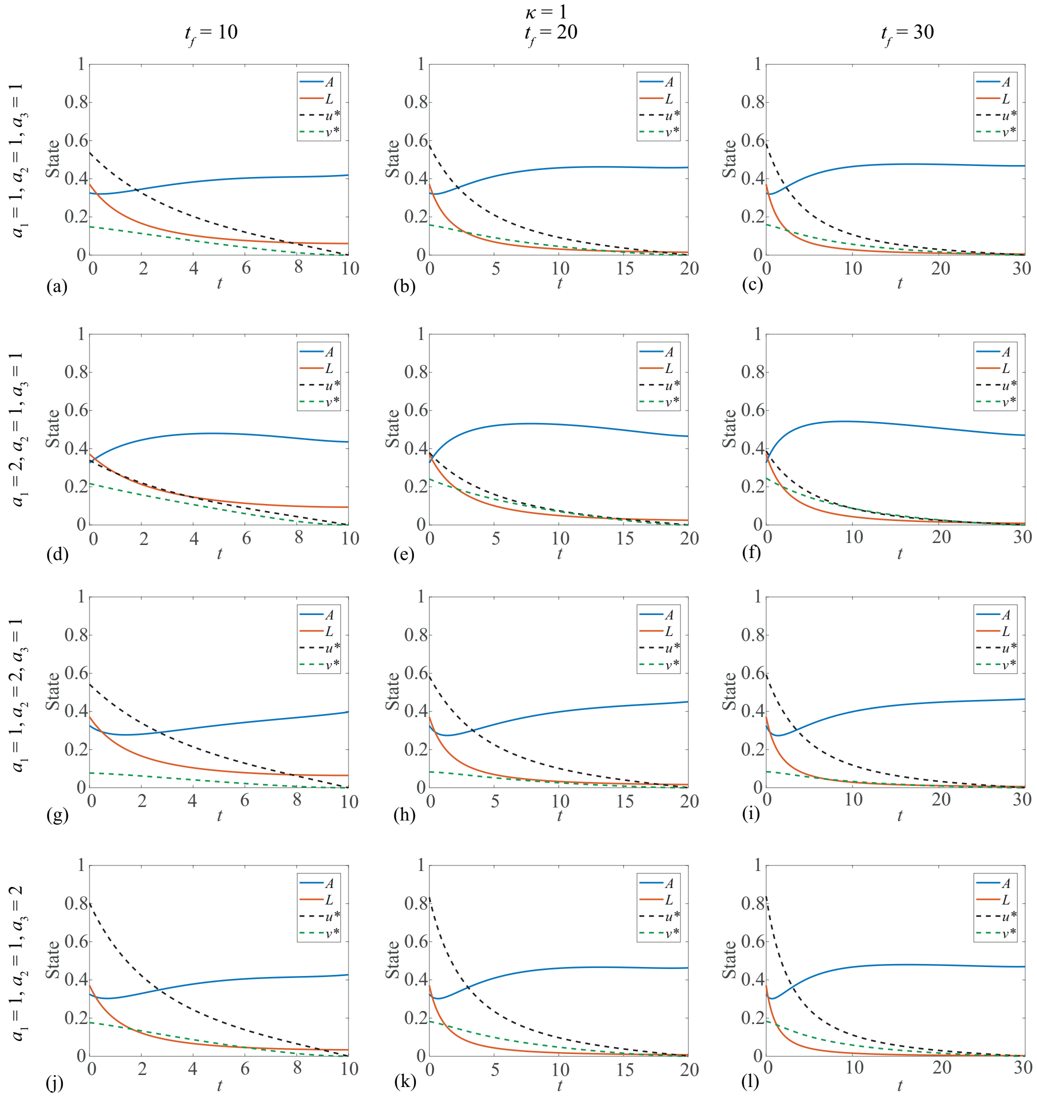


Fig. S2. Results are presented where both controls applied continuously, under a range of pay-off weightings ($a_1, a_2, a_3 \in \{1, 2\}$) and where each column of the figure corresponds to a different terminal time; $t_f \in \{10, 20, 30\}$. This figure corresponds to the pay-off given by Equation (S2), where the leukaemic term enters the pay-off linearly. This corresponds to Equation (4) in the Main document with $p = 2$, $q = 2$, $r = 1$.

##### 1.2 *Continuous chemotherapy control, bang-bang stem cell transplant control*

This Subsection contains results for pay-offs presented in Equation (S3) and Equation (S4), where the chemotherapy control is continuous and the stem cell transplant control is bang-bang. We also apply bounds to the continuous chemotherapy control. These pay-offs correspond to results in Figure S3 and Figure S4, respectively. The pay-off functions are given by:

$$J = \int_0^{t_f} (a_1 u^2 + a_2 v + a_3 L^2)\ \mathrm{d}t, \quad 0 \leq u \leq u_b, \quad 0 \leq v \leq v_b, \tag{S3}$$

$$J = \int_0^{t_f} (a_1 u^2 + a_2 v + a_3 L)\ \mathrm{d}t, \quad 0 \leq u \leq u_b, \quad 0 \leq v \leq v_b. \tag{S4}$$

When the stem cell control is bang-bang, we only observe it switching on when its weighting in the pay-off is significantly lower than the weightings for the chemotherapy control and the leukaemia. Primarily, this is because the pay-off only considers the leukaemic population (and not the progenitor blood cell population), and the stem cell control is not as effective at reducing the leukaemic population as it can only achieve this indirectly through the competition between progenitor blood and leukaemia. In addition, since the bang-bang control arises from a linear term in the pay-off, it is more costly than an equivalently weighted quadratic component applied at the same level.

Bounded continuous controls allow us to incorporate physical constraints such as a maximum tolerable dose. In Figure S3 an upper bound of $u_b = 0.3$ is placed on the continuous control. This leads to interesting results, particularly in combination with order of magnitude variations in $\kappa$. When all pay-off terms are weighted equally, as in Figure S3(a-c), increasing $\kappa$ increases the duration of the interval where chemotherapy is applied at its upper bound. Though initially counter-intuitive, this can be explained by the competition between leukaemia and progenitor blood cells; as $\kappa$ is increased the progenitor population is further reduced and therefore presents weaker competition to the leukaemia.

With large $\kappa$ and a low weighting on leukaemia in the pay-off relative to the controls, as in Figure S4(l), we observe that only a small amount of chemotherapy control is applied, near the terminal time. This is due to a combination of not incorporating progenitor cells directly in the pay-off and having a fixed terminal time. As the chemotherapy control is quadratic in the pay-off, the small amount of chemotherapy control applied would contribute very little to the pay-off, such that the resulting minor reduction in leukaemia is worthwhile. However, this is only worthwhile near the end of the time interval. The chemotherapy control is not applied earlier in the interval as the cumulative effect of reduced competition due to decreased progenitor population would outweigh the benefit of applying the chemotherapy. Similar behaviour appears in Figure S4(i), corresponding to large $\kappa$ and a low weighting on the stem cell transplant control in the pay-off.

We investigate the impact of varying the upper bound on the continuous control in Figure S4. As the upper bound is increased, the interval over which the continuous control is applied at its upper bound is reduced. Adjusting the upper bound produces a different response from the state, and therefore also alters the control dynamics over the intervals where the control is applied at a lower level than the upper bound. It follows that when the control is never applied at the upper bound (Figure S4(j,k,l)), changing the upper bound does not impact the dynamics of the system.

Comparing Figure S4(h) with Figure S3(h), we see that incorporating leukaemia in the pay-off linearly rather than quadratically increases the interval over which the bounded continuous chemotherapy control is applied at its upper bound. The bang-bang stem cell transplant control is applied for longer.

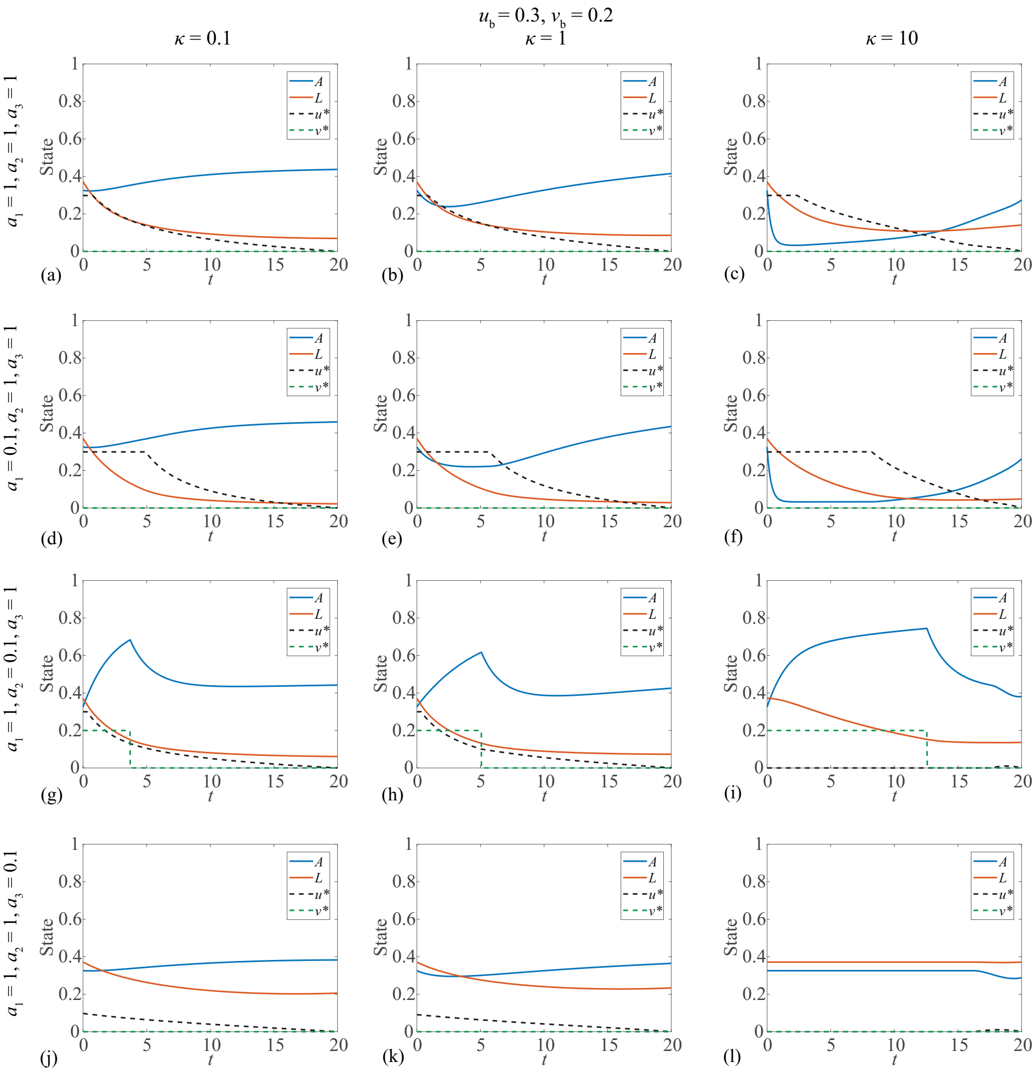


Fig. S3. Results are presented for a bounded continuous chemotherapy control ($u_b = 0.3$), and bang-bang stem cell transplant control ($v_b = 0.2$). Results correspond to a range of pay-off weightings ($a_1, a_2, a_3 \in \{0.1, 1\}$) and examine a more extreme variation of $\kappa$ than in Figure S1. This figure corresponds to the pay-off given by Equation (S3), where the leukaemic term enters the pay-off quadratically. This corresponds to Equation (4) in the Main document with $p = 2$, $q = 1$, $r = 2$.

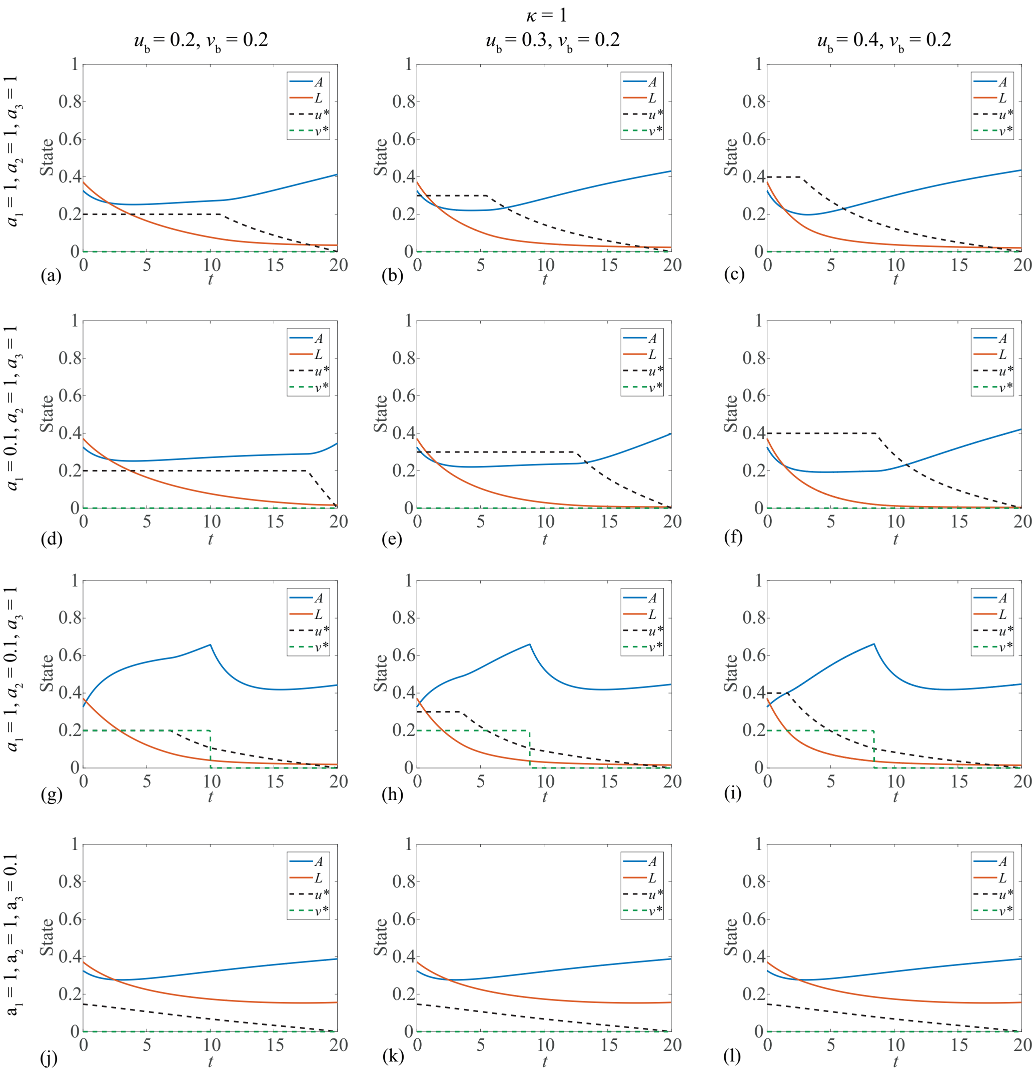


Fig. S4. Results are presented for a bounded continuous chemotherapy control, and bang-bang stem cell transplant control ($v_b = 0.2$). The upper bound on the continuous chemotherapy control is varied; $u_b \in \{0.2, 0.3, 0.4\}$, under a range of pay-off weightings ($a_1, a_2, a_3 \in \{0.1, 1\}$). This figure corresponds to the pay-off given by Equation (S4), where the leukaemic term enters the pay-off linearly. This corresponds to Equation (4) in the Main document with $p = 2$, $q = 1$, $r = 1$.

##### 1.3 *Bang-bang chemotherapy control, continuous stem cell transplant control*

This Subsection contains results for pay-offs presented in Equation (S5) and Equation (S6), where the chemotherapy control is bang-bang and the stem cell transplant control is continuous. These pay-offs correspond to results in Figure S5 and Figure S6, respectively. The pay-off functions are given by:

$$J = \int_0^{t_f} (a_1 u + a_2 v^2 + a_3 L^2)\ \mathrm{d}t, \quad 0 \le u \le u_b, \tag{S5}$$

$$J = \int_0^{t_f} (a_1 u + a_2 v^2 + a_3 L)\ \mathrm{d}t. \quad 0 \le u \le u_b. \tag{S6}$$

We revisit varying the terminal time in Figure S5 to investigate its impact on the dynamics of bang-bang controls. The dynamics are again similar with different terminal times, but are not direct scalings. As the terminal time increases, the bang-bang control is applied over a longer interval, although this interval does not necessarily increase proportionally with the terminal time. This is clear in the second row of Figure S5; where the bang-bang control switches off around $t = 8$ when $t_f = 10$, and switches off around $t = 15$ when $t_f = 30$.

When the continuous stem cell transplant control is weighted much lower than the other terms in the pay-off, it is applied liberally (S5(g,h,i)). In this situation the bang-bang chemotherapy control is not switched on, as the leukaemic population is reduced through the competition between $A$ and $L$. We note that this level of stem cell transplant control is not physically realistic, as it increases the progenitor population well above the carrying capacity of the model, resulting in a significant decline in the progenitor population as the support from the stem cell transplant control is reduced. These kind of results can provide insight regarding the quality of the model and/or the pay-off form and weightings. If we believe the model to be sufficiently realistic, then obtaining non-physical control results suggests that the pay-off form or weightings must not be realistic; in this case it would appear that applying the stem cell transplant control is not costly enough.

In Figure S6, we present results exploring the continuous stem cell control and bang-bang chemotherapy control dynamics with leukaemia entering the pay-off linearly. We consider the case where the continuous stem cell transplant control is unbounded, and also consider imposing upper bounds of $v_b = 0.1$ and $v_b = 0.3$. When the stem cell transplant control is weighted lower in the pay-off, similarly to Figure S5(g,h,i), the continuous stem cell transplant control is applied too liberally. This is partially mitigated through imposing the upper bounds, as in Figure S6(g,i), although we still see a sharp decline in the progenitor cell population once the support from the control is removed. This highlights a challenge in determining appropriate weightings of multiple controls with different forms; the form of the control terms in the pay-off (linear, quadratic) directly impacts their contribution to the pay-off. Pay-off terms are equally weighted in Figure S6(a), and the continuous stem cell control is applied at the upper bound of $v_b = 0.1$, matching the bound of the chemotherapy control at $u_b = 0.1$. The contribution of the chemotherapy control to the pay-off is an order of magnitude greater than that of the stem cell control.

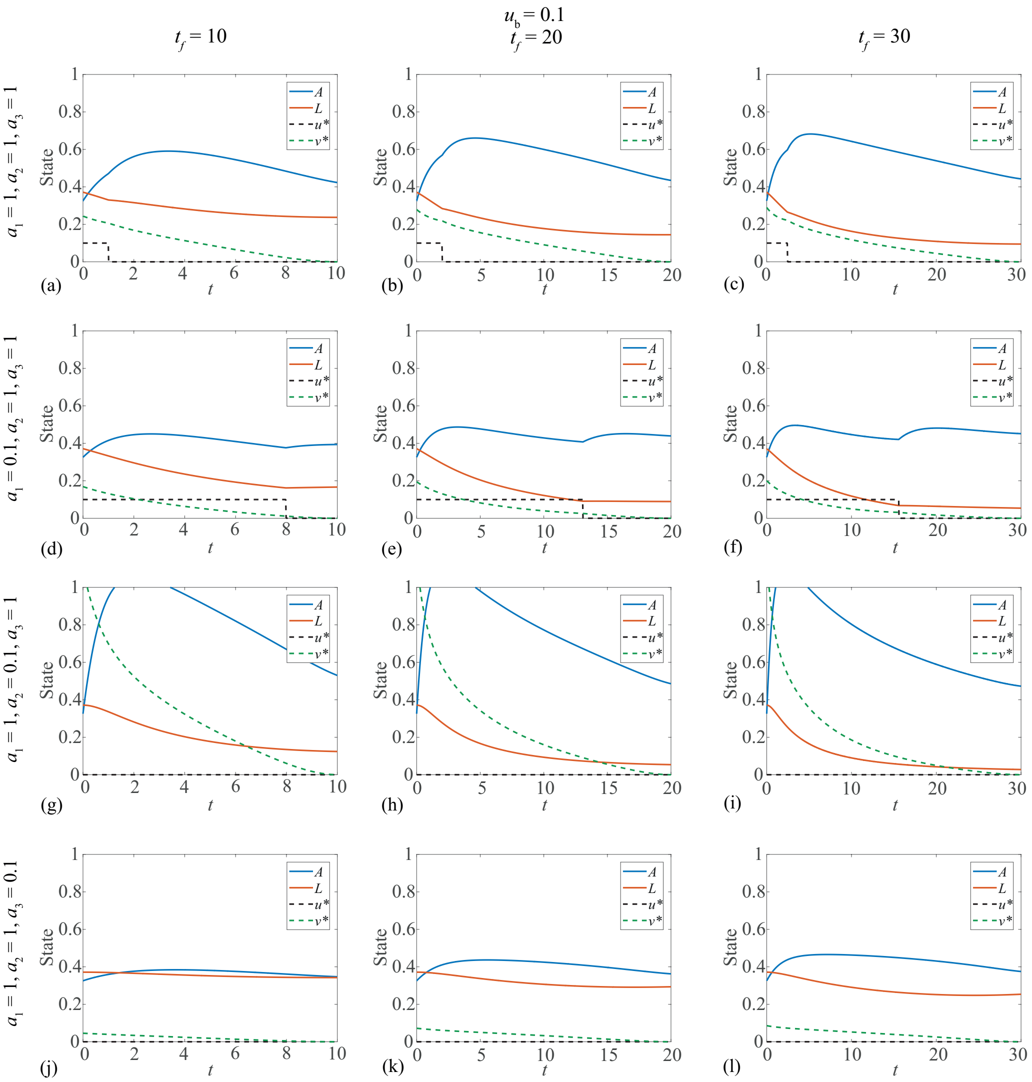


Fig. S5. Results are presented for a bang-bang chemotherapy control and continuous stem cell transplant control, with an upper control bound of 0.1 on the bang-bang control. Each column of the figure corresponds to a different terminal time; ($t_f \in \{10, 20, 30\}$). This figure corresponds to the pay-off given by Equation (S5), where the leukaemic term enters the pay-off quadratically. This corresponds to Equation (4) in the Main document with $p = 1$, $q = 2$, $r = 2$.

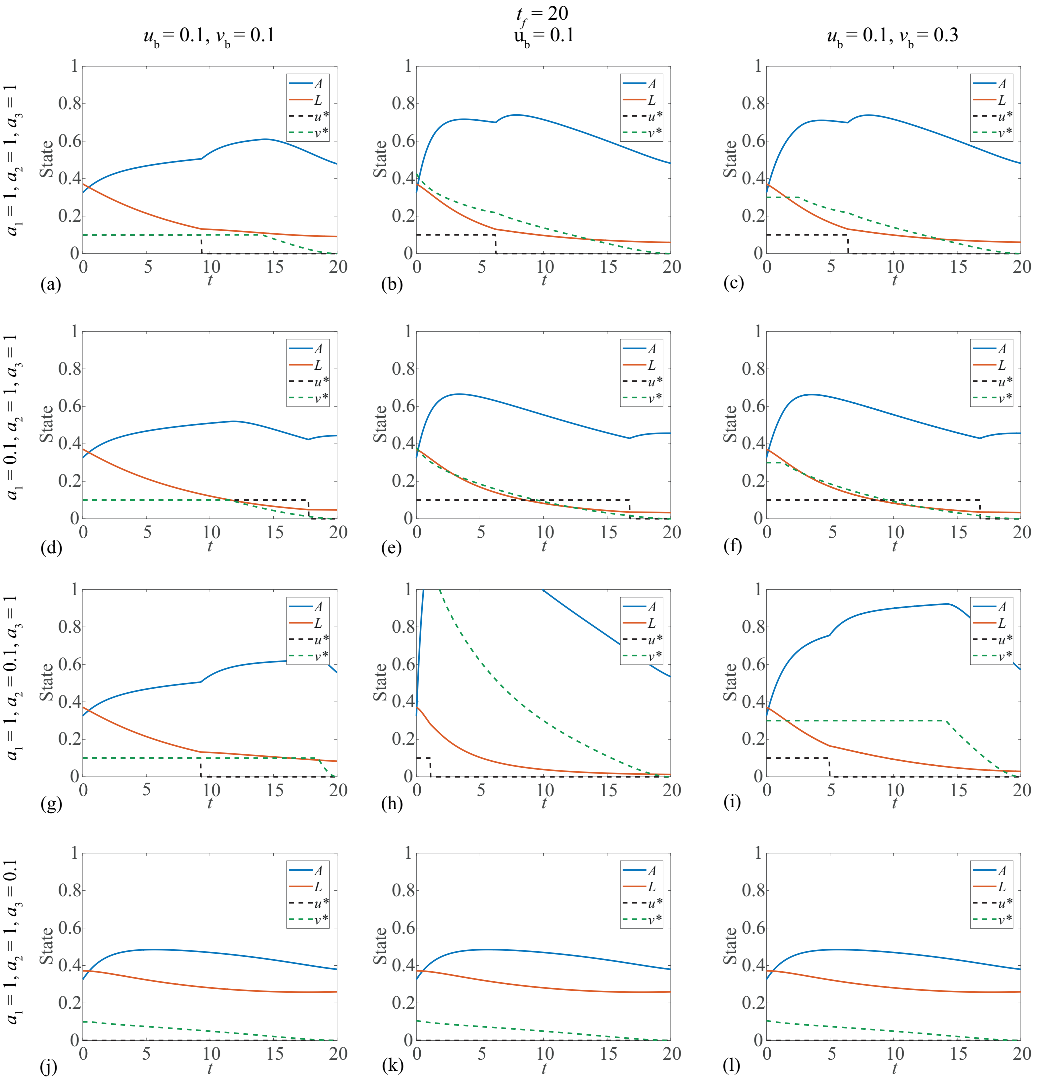


Fig. S6. Results are presented for a bang-bang chemotherapy control ($u_b = 0.1$) and continuous stem cell transplant control, under a range of pay-off weightings ($a_1, a_2, a_3 \in \{0.1, 1\}$). The continuous stem cell transplant control is not constrained by an upper bound in the central column, with an upper bound of $v_b = 0.1$ in the left column and $v_b = 0.3$ in the right column. This figure corresponds to the pay-off given by Equation (S6), where the leukaemic term enters the pay-off linearly. This corresponds to Equation (4) in the Main document with $p = 1$, $q = 2$, $r = 1$.

##### *1.4 Bang-bang chemotherapy control, bang-bang stem cell transplant control*

This Subection contains results for pay-offs presented in Equation (S7) and Equation (S8), where both controls are bang-bang. These pay-offs correspond to results in Figure S7 and Figure S8, respectively. The pay-off functions are given by:

$$J = \int_0^{t_f} (a_1 u + a_2 v + a_3 L^2)\ \mathrm{d}t, \quad 0 \le u \le u_b, \quad 0 \le v \le v_b, \tag{S7}$$

$$J = \int_0^{t_f} (a_1 u + a_2 v + a_3 L)\ \mathrm{d}t, \quad 0 \le u \le u_b, \quad 0 \le v \le v_b. \tag{S8}$$

The intuitive results that we observe in previous subsections are reflected in Figure S7 with both controls bang-bang. In the central column we consider bang-bang controls with equal upper bounds: $u_b = v_b = 0.1$. In the left column, we increase the upper bound on the stem cell transplant control to $v_b = 0.2$ and in the right column we increase the upper bound on the chemotherapy control to $u_b = 0.2$. This enables us to investigate the impact of control strength. As the upper bound on the controls increase, the duration that they are switched on decreases.

When all pay-off terms are equally weighed, only the chemotherapy control is applied, as it is more effective at reducing the leukaemic population. When the pay-off weighting of a bang-bang control is reduced, it is applied for longer. Increasing the upper bound on one control can reduce the duration that the other control is required, for example compare Figure S7(h) with Figure S7(i). With both controls bang-bang and all terms weighed equally, the final populations indicate that the leukaemia will return to its coexisting steady state; such a result suggests that the leukaemia may not be sufficiently weighted in the pay-off.

In Figure S8 we investigate the impact of varying $\kappa$ when the controls are bang-bang, and when leukaemia enters the pay-off linearly. For the small variations considered in this case ($\kappa \in 0.5, 1, 2$) we notice changes in the state response, particularly of the progenitor population, but change to the control dynamics is marginal. As $\kappa$ is increased, leading to a greater reduction in the progenitor population; the stem cell transplant control is applied for longer, and the chemotherapy control is applied for less time.

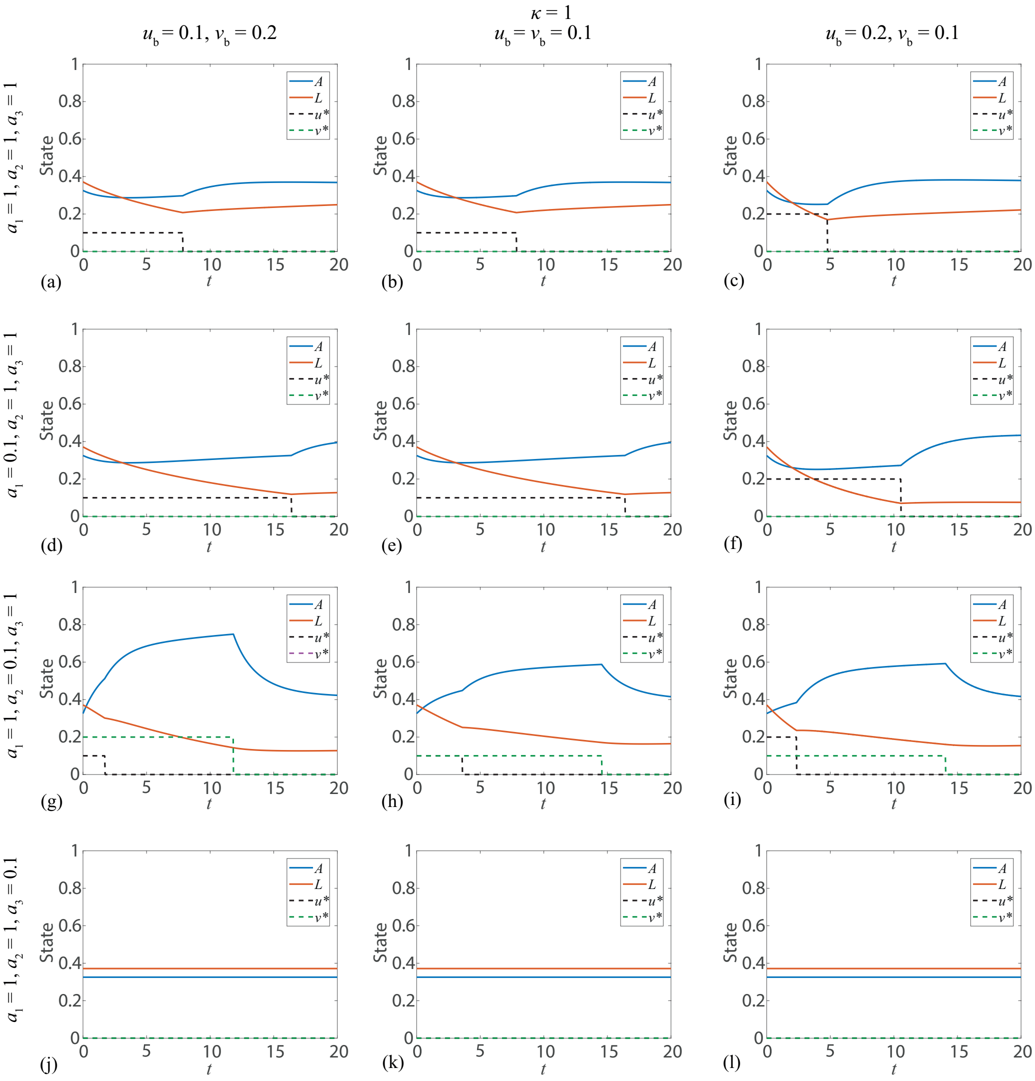


Fig. S7. Results are presented where both controls are bang-bang, under a range of pay-off weightings ($a_1, a_2, a_3 \in \{0.1, 1\}$). In the central column we consider equal control bounds; $u_b = v_b = 0.1$. In the left column the upper bound on the stem cell transplant control is increased ($v_b = 0.2$), and in the right column the upper bound on the chemotherapy control is increased ($u_b = 0.2$). This figure corresponds to pay-off given by Equation (S7), where the leukaemic term enters the pay-off quadratically. This corresponds to Equation (4) in the Main document with $p = 1$, $q = 1$, $r = 2$.

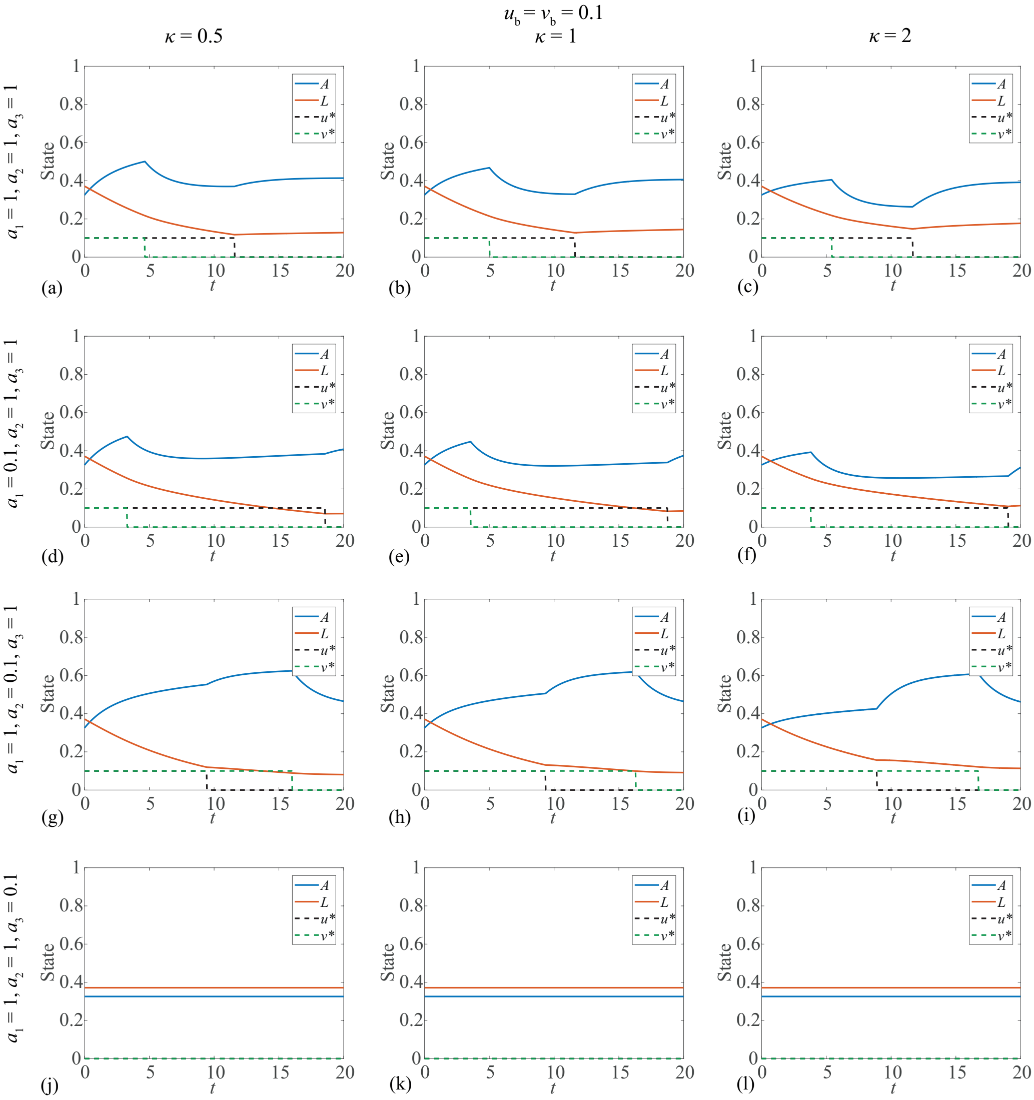


Fig. S8. Results are presented where both controls are bang-bang, under a range of pay-off weightings ($a_1, a_2, a_3 \in \{0.1, 1\}$) and for upper control bounds $u_b = v_b = 0.1$. We investigate how $\kappa$, the rate that chemotherapy kills progenitor blood cells relative to leukaemic cells, impacts the dynamics of bang-bang controls. This figure corresponds to the pay-off given by Equation (S8), where the leukaemic term enters the pay-off linearly. This corresponds to Equation (4) in the Main document with $p = 1$, $q = 1$, $r = 1$.

#### 2 Supporting code

Code for implementation of the optimal control algorithms in this work is made freely available on GitHub.

#### 3 Forward backward sweep method

In Algorithm 1 we present a concise algorithm for the FBSM with multiple controls [2]. We denote the system state as $\mathbf{x}(t)$, the co-state as $\boldsymbol{\lambda}(t)$, and the controls as $\mathbf{u}(t)$. For the two control model considered in this work, $\mathbf{x}(t) = [A(t), L(t)]^{\mathrm{T}}$, $\mathbf{u}(t) = [u(t), v(t)]^{\mathrm{T}}$, and $\boldsymbol{\lambda}(t) = [\lambda_1(t), \lambda_2(t)]^{\mathrm{T}}$; where $\lambda_1(t)$ and $\lambda_2(t)$ are the co-state equations derived from the Hamiltonian [2].

**Algorithm 1: FBSM for multiple controls**

i. Make an initial guess of $\mathbf{u}(t)$.
   *For all problems considered in this work, $u(t) \equiv v(t) \equiv 0$ is sufficient.*
ii. Using the initial condition $\mathbf{x}(0) = \mathbf{x}_0$, solve for $\mathbf{x}(t)$ forward in time using the initial guess of $\mathbf{u}(t)$.
iii. Using the transversality condition $\boldsymbol{\lambda}(t_f)$, solve for $\boldsymbol{\lambda}(t)$ backwards in time, using the values for $\mathbf{u}(t)$ and $\mathbf{x}(t)$.
iv. Calculate $\mathbf{u}_{\mathrm{new}}(t)$ by evaluating the expression for the optimal control $\mathbf{u}^*(t)$ using the updated $\mathbf{x}(t)$ and $\boldsymbol{\lambda}(t)$ values.
v. Update $\mathbf{u}(t)$ based on a combination of $\mathbf{u}_{\mathrm{new}}(t)$ and the previous $\mathbf{u}(t)$.
vi. Check for convergence.
   *If $\mathbf{x}(t)$, $\boldsymbol{\lambda}(t)$ and $\mathbf{u}(t)$ meet a specified tolerance, accept $\mathbf{x}(t)$, $\boldsymbol{\lambda}(t)$ and $\mathbf{u}(t)$, otherwise return to Step* ii. *using the updated $\mathbf{u}(t)$.*
